# Canonical cortical architecture supports the emergence of noise-invariant auditory representations

**DOI:** 10.64898/2025.12.19.695511

**Authors:** Tomas Suarez Omedas, Ross S. Williamson

**Affiliations:** Neuroscience Institute, Carnegie Mellon University, Pittsburgh, PA; Department of Otolaryngology, University of Pittsburgh, Pittsburgh, PA; Department of Neurobiology, University of Pittsburgh, Pittsburgh, PA; Department of Bioengineering, University of Pittsburgh, Pittsburgh, PA; Pittsburgh Hearing Research Center, University of Pittsburgh, Pittsburgh, PA; Center for the Neural Basis of Cognition, University of Pittsburgh, Pittsburgh, PA

## Abstract

Neurons in the auditory system must represent behaviorally relevant sounds in the presence of background noise (BN) to support noise-invariant perception and behavior. Although primary auditory cortex (ACtx) has been implicated in constructing noise-invariant representations, it remains unclear which excitatory subpopulations within ACtx carry out this transformation from noise-dependent to noise-invariant coding. To address this, we presented pure tones with and without continuous BN to head-fixed mice and used two-photon calcium imaging to record sound-evoked activity from three major excitatory subpopulations in ACtx: layer (L)2/3 intratelencephalic (IT) neurons, L5 IT neurons, and L5 extratelencephalic (ET) neurons. L2/3 IT neurons exhibited strong noise dependence at the level of single-neuron responses, pairwise interactions, and population representations. In contrast, deep-layer pathways showed greater noise invariance, with L5 IT neurons preserving stable representations most consistently and L5 ET neurons exhibiting more limited invariance at the population level. These findings reveal a functional division of labor in ACtx, in which superficial neurons remain noise-dependent and deep-layer broadcast pathways, particularly L5 IT, preferentially carry noise-invariant representations, suggesting that excitatory subpopulations contribute differentially to the construction and propagation of noise-invariant codes.

## Introduction

The auditory system continuously faces the task of accurately representing sound features, even if they are embedded in background noise (BN). Robust, or ideally invariant, coding of acoustic attributes such as frequency and intensity enables animals to detect threats and allows humans to understand speech in complex acoustic scenes (e.g., the cocktail-party setting). The construction of such a noise-invariant neural code, one that represents acoustic characteristics regardless of BN, poses a central challenge for auditory processing. As BN increases and the signal-to-noise ratio (SNR) worsens, the acoustic waveform at the ear is distorted, cochlear responses are reshaped, and these changes propagate through the ascending auditory pathway [1–9]. Despite extensive work on BN effects across numerous auditory brain regions [2, 10–14], the precise locus and circuit mechanisms by which noise-invariant representations first emerge remain unresolved.

Sounds reaching the ear are decomposed by the cochlea into frequency components that are relayed along the ascending auditory pathway [15, 16]. When sounds are embedded in BN, their spectrotemporal structure becomes distorted, producing measurable changes in neural representations in the auditory periphery [10, 11], midbrain [12], and cortex [2, 13, 14]. BN can modify both single-neuron and population responses, shifting firing rates and reshaping population activity patterns [10–12], degrading spatial cues [10], suppressing sound-evoked activity [2, 17], and modulating baseline firing [1]. Across the pathway, representations generally become more noise-invariant as signals ascend toward auditory cortex (ACtx)[1, 9]; responses in primary ACtx are closer to noise-invariant than in the periphery yet remain measurably affected by BN [4, 18], whereas non-primary fields often exhibit stronger invariance [4, 9, 19]. These observations identify ACtx as a critical locus where invariance to BN is refined, motivating the question of how cortical circuits support this transformation.

ACtx is a hub for sound processing and the brain-wide broadcast of auditory information, roles that are particularly important in complex acoustic environments where sounds are embedded in BN [7]. Inactivation of ACtx significantly impairs pure tone detection in BN, and disrupting somatostatin- or parvalbumin-positive interneurons produces similar behavioral deficits [6, 20], underscoring a role for ACtx in supporting noise-invariant perception. Building on these findings, recent work suggests that ACtx transforms noise-dependent inputs into noise-invariant representations through adjustments of excitatory-inhibitory balance [2, 5, 6] and cholinergic neuromodulation [3]. Nevertheless, the computations that implement noise invariance remain unclear, in part because prior studies have not considered the functional diversity of excitatory subpopulations within the cortical microcircuit.

Within ACtx, excitatory neurons are stratified across layers and projection classes [21–26]. Two major subpopulations, intratelencephalic (IT) and extratelencephalic (ET) neurons, differ in their anatomical, genetic, and functional properties [27–29]. In ACtx, layer (L)2/3 IT neurons form dense intracortical networks and are positioned to perform local computations on incoming sensory input [24]. In contrast, L5 IT and ET neurons constitute broadcast pathways: L5 IT neurons project widely within the telencephalon [26, 28, 30–33], whereas L5 ET neurons target subcortical structures such as the inferior colliculus and thalamus [25, 26, 29]. These subpopulations make distinct contributions to cortical function, including learning perceptually relevant information [22], amplifying sensory signals after injury [32], and regulating thalamic transmission [25]. Given these divergent roles and projection patterns, we tested whether L2/3 neurons are more susceptible to BN, and whether L5 IT and L5 ET neurons preferentially transmit noise-invariant signals to downstream targets.

We examined noise invariance at three analytical levels (single-neuron, pairwise, and population) in three major ACtx excitatory subpopulations: L2/3, L5 IT, and L5 ET. Using *in vivo* two-photon calcium imaging in mice, we recorded responses to pure tones presented with and without BN. L2/3 neurons exhibited pronounced noise-dependence at all three levels, while both both L5 IT and L5 ET populations exhibited robust, noise-invariant coding. Thus, while superficial neurons alter their representations in the presence of noise, deep-layer output pathways maintain stable representations, indicating that noise invariance is preferentially expressed within the cortical output stages of ACtx.

## Results

To assess noise invariance in sound representations across excitatory subpopulations of ACtx, we performed *in vivo* two-photon calcium imaging to record activity from three targeted groups: L2/3 (*n* = 961 neurons; *N* = 7 mice), L5 IT (*n* = 2576; *N* = 6) and L5 ET (*n* = 566; *N* = 4) (Fig 1A). GCaMP8s expression was selectively driven in each subpopulation using genetic and viral strategies (see Methods). Head-fixed mice were imaged through a cranial window over right ACtx, while pure tones were delivered monaurally to the left ear (Fig 1B). Tones spanned 4–45 kHz in half-octave steps and 20–70 dB SPL in 10 dB SPL steps (8 frequencies × 6 intensities = 48 unique conditions), each repeated 20 times. Each mouse completed two sessions: one without BN (No-BN) and one with continuous BN at 50 db SPL (Fig 1C–D). Fluorescence calcium traces were extracted and deconvolved using standard approaches [34, 35]. All deconvolved responses were z-scored relative to their pre-stimulus baseline (Fig 1E).

**Figure 1:**
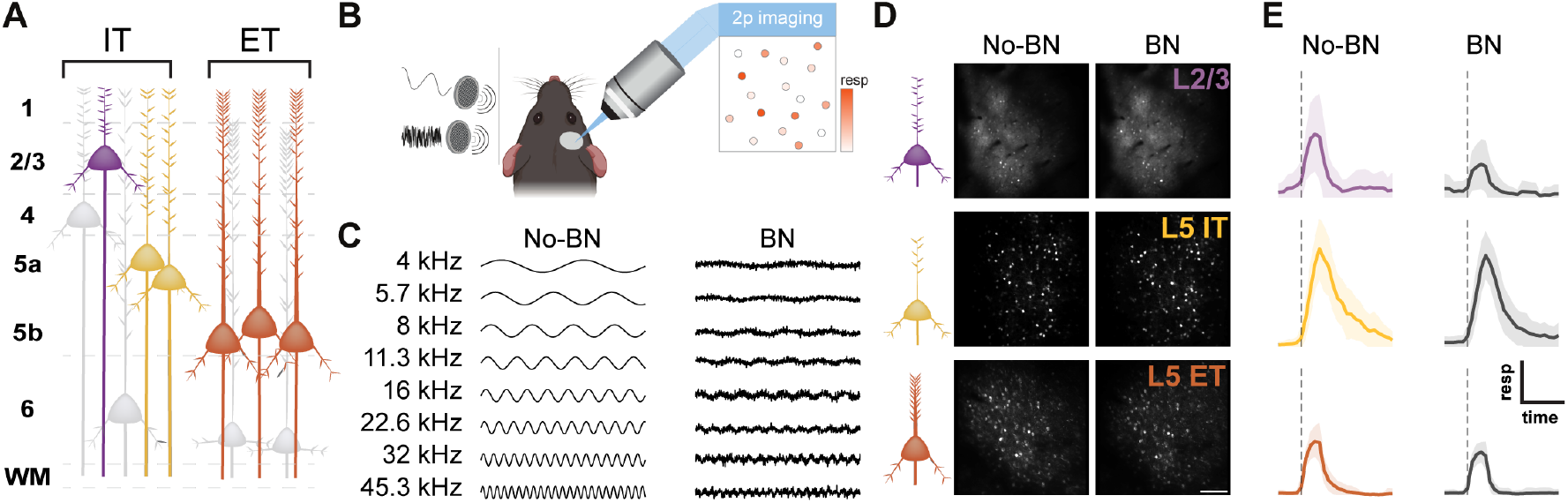
Two-photon recordings from awake, head-fixed mice. **(A)** Schematic illustrating excitatory subpopulations in auditory cortex (ACtx). **(B)** Two-photon imaging setup for recordings in awake, head-fixed mice. **(C)** Schematic of acoustic stimulation in the absence (No-BN) and presence (BN) of continuous background noise. **(D)** Example fields of view (FoVs) from each recorded subpopulation. Imaging depths were 206 *µ*m (L2/3), 415 *µ*m (L5 IT), and 494 *µ*m (L5 ET). **(E)** Example sound-evoked calcium responses at the best frequency for one representative neuron from each FoV shown in **D**.

### BN attenuates single-neuron responses in L2/3

We first asked whether single-neuron responses in each excitatory subpopulation are noise-invariant. To do so, we analyzed two complementary aspects of single-neuron activity: (i) mean stimulus-locked responses, summarized by frequency and intensity tuning curves, and (ii) trial-to-trial response distributions, which capture variability not reflected in the mean. Tuning analyses assessed additive and multiplicative changes between No-BN and BN conditions, whereas distributional analyses quantified stimulus information and its modulation by BN.

For each sound-responsive neuron (see Methods), we computed frequency response areas (FRAs) by averaging sound-evoked activity across all trials for each frequency–intensity combination (Fig 2A). From these FRAs, we derived frequency and intensity tuning curves for the No-BN and BN conditions by marginzalizing over the complementary stimulus dimension. To compare frequency tuning across noise conditions, we aligned each neuron’s tuning curve to its best frequency (BF) and averaged the resultant BF-centered curves for each subpopulation. BF-centered frequency tuning curves showed a significant reduction in L2/3, but no change in L5 IT or ET (Fig 2B top row, two-way ANOVA, main effect for BN, L2/3: *p* = 0.0135; L5 IT: *p* = 0.9014; L5 ET: *p* = 0.8156). This reduction was accompanied by a significant interaction between frequency and BN in L2/3, which is concentrated at best frequency (interaction between frequency and BN: *p* = 2.5946 × 10^−4^; Tukey-Kramer post hoc test for best frequency: *p* = 5.1255 × 10^−13^). Average intensity tuning curves likewise decreased in L2/3 but not in L5 IT or ET (Fig 2B bottom row, two-way ANOVA, main effect for BN, L2/3: *p* = 3.1543 × 10^−12^, L5 IT: *p* = 0.8450, L5 ET: *p* = 0.8609). Together, these results indicate that, on average, L2/3 neurons tuning curves are suppressed in the presence of BN, while L5 IT and L5 ET curves remain unchanged.

**Figure 2:**
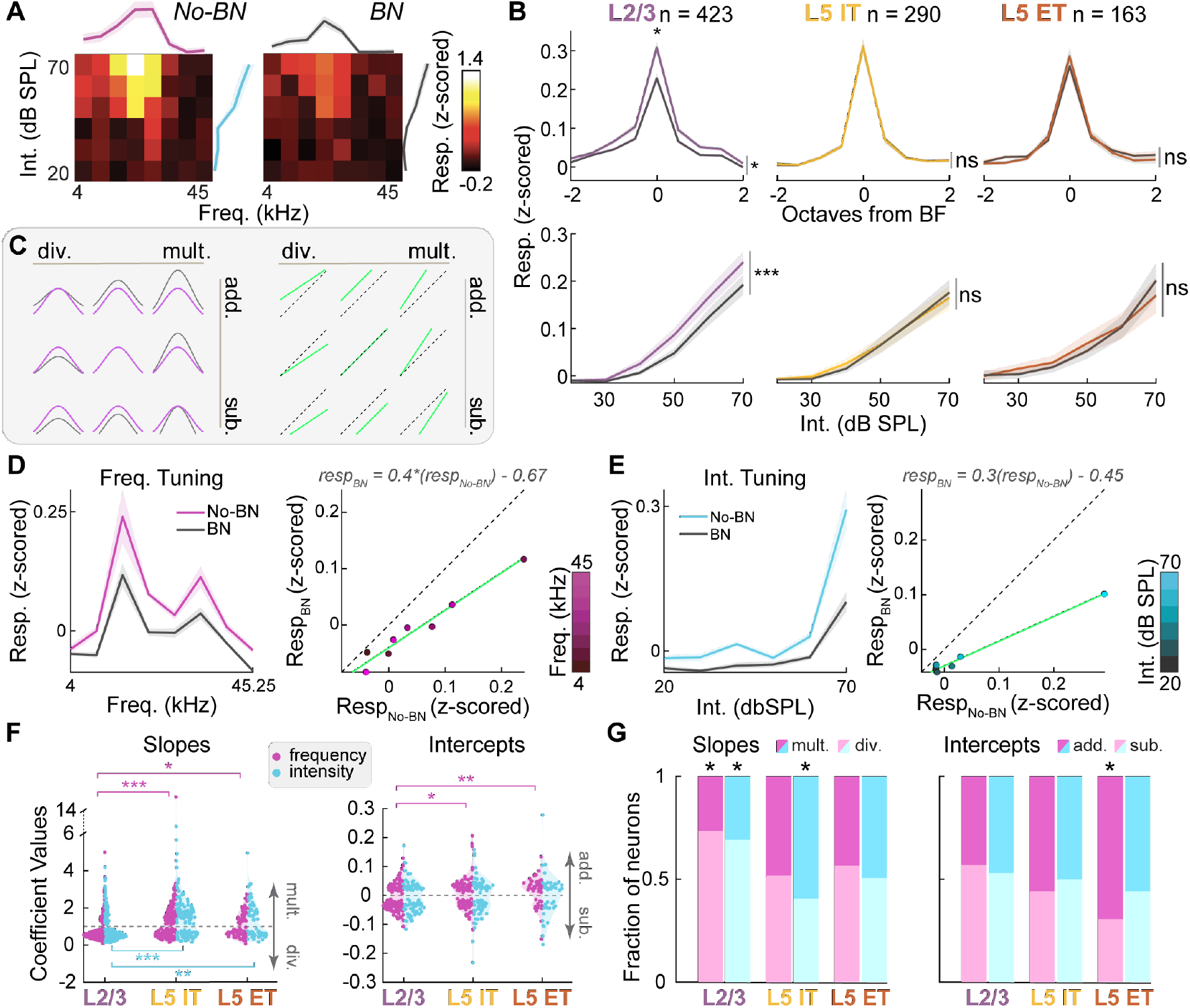
ACtx broadcast subpopulations exhibit noise-invariant single-neuron responses. **(A)** Example frequency response area (FRA) from an L2/3 neuron in the absence (No-BN, left) and presence (BN, right) of background noise. Both panels use the same color scale, with lighter colors indicating larger responses. **(B)** Average best-frequency-centered frequency tuning curves (top row) and average intensity tuning curves (bottom row) for each excitatory sub-population. **(C)** Schematic illustrating linear transformations between tuning curves across BN conditions (left) and reduced major axis (RMA) regression used to quantify these changes (right). Dotted line indicates the unity line. **(D)** Example frequency tuning curves for a single neuron recorded in No-BN and BN (left) and corresponding RMA regression (right). Estimated slope and intercept coefficients are shown above the plot. **(E)** Same as **D**, but for intensity tuning curves. **(F)** Distributions of RMA slope coefficients (left) and intercept coefficients (right) that differed significantly from 1 and 0, respectively, for frequency (magenta) and intensity (cyan) tuning curves. Number of neurons with significant regressions for L2/3, L5 IT and L5 ET respectively: *n* = 272, 204, 88 frequency slopes, *n* = 222, 140, 65 intensity slopes, *n* = 161, 113, 49 frequency intercepts, *n* = 107, 82, 43 intensity intercepts. **(G)** Proportions of RMA coefficients classified as additive, subtractive, multiplicative, or divisive for each subpopulation. Same *n* as in **F**.

Although averaged tuning curves showed that L2/3 responses were altered differently from those in L5 IT and L5 ET, these averages do not capture how individual neurons adjusted their tuning curves in BN. To quantify neuron-by-neuron changes that underlie the population-level trends, we performed reduced major axis (RMA) regression between each neuron’s No-BN and BN tuning curves (Fig 2C) [32, 36]. This approach estimates how much a tuning curve shifts (additive/subtractive) and/or scales (multiplicative/divisive) in BN (Fig 2C). Each neuron produced four coefficients: a slope and intercept for its frequency tuning curve, and a slope and intercept for its intensity tuning curve (Fig 2D-E). Slope values greater than one indicated multiplicative scaling, whereas values less than one indicated divisive scaling. Similarly, intercept coefficients greater than zero indicated additive shifts, whereas values less than zero indicated subtractive shifts.

The distribution of slope coefficients differed across subpopulations (Fig 2F left, Kruskal–Wallis test, main effect for frequency slope: *p* = 2.5484 × 10^−6^; main effect for intensity slope: *p* = 5.3103 × 10^−7^). L2/3 neurons showed lower slope values than both L5 IT and ET populations for frequency and intensity tuning (Dunn–Šidák post hoc test for frequency and intensity curves, respectively, L2/3-L5 IT: *p* = 3.1471 × 10^−6^ and *p* = 1.8054 × 10^−6^, L2/3-L5 ET: *p* = 0.0114 and *p* = 0.0021, L5 IT-L5 ET: *p* = 0.8289 and *p* = 0.9683), indicating that BN produced a stronger divisive scaling of responses in L2/3 than in either deep-layer subpopulation.

Intercept coefficients for frequency tuning curves also differed across subpopulations (Fig 2F right, Kruskal-Wallis test, main effect for frequency intercept: *p* = 0.0020), with L2/3 neurons showing lower intercepts than L5 IT and L5 ET neurons (Dunn–Šidák post hoc test, L2/3-L5 IT *p* = 0.0304, L2/3-L5 ET: *p* = 0.0058, L5 IT-L5 ET: *p* = 0.6027). However, no measurable differences in intensity tuning curve intercepts were recorded (Fig 2F, right, Kruskal–Wallis test, main effect for intensity intercept: *p* = 0.6599).

To assess overall trends in tuning curve modulation across subpopulations, we computed, for each subpopulation, the proportion of neurons showing suppressive versus enhancing changes. Deviations above or below 50% indicate whether a subpopulation tends to increase or decrease its responses in BN. If the effect were not subpopulation-specific, these proportions would be similar across subpopulations.

We found that the proportions of multiplicative versus divisive coefficients differed between subpopulations for both frequency and intensity slopes (Fig 2G left, separate chi-square tests for frequency and intensity slopes, respectively; *p* = 3.5892 × 10^−6^ and *p* = 2.971 × 10^−7^), as well as for additive versus subtractive frequency intercepts (Fig 2G right, chi-square test for frequency intercepts: *p* = 0.0026). In L2/3 neurons, the proportions of frequency and intensity slopes were strongly biased toward divisive changes (binomial test for frequency and intensity slopes: *p* = 4.2188 × 10^−15^ and *p* = 7.6385 × 10^−9^). In contrast, L5 IT and L5 ET neurons showed proportions closer to 50%, with the only significant biases favoring enhancement of tuning curves via multiplicative or additive changes, specifically in the intensity slopes of L5 IT neurons (binomial test: *p* = 0.0342) and in the frequency intercepts of L5 ET neurons (binomial test: *p* = 0.0094). Together, these results indicate that, although individual neurons in all subpopulations can either enhance or suppress their tuning, L2/3 neurons tended to exhibit predominantly divisive changes, whereas L5 IT and ET neurons maintain more balanced, and in some cases mildly enhancing, tuning shifts.

While changes in tuning curves reveal the direction of modulation induced by BN, they do not capture whether a neuron’s trial-to-trial response distribution changes under BN. For example, two neurons may show similar suppression in mean tuning across conditions, yet differ in how reliably they respond to each tone. Such differences in response variability reflect how stable a neuron’s responses remain after BN-induced changes in mean tuning. To quantify this, we applied an information-theoretic framework to measure how much of the variability in each neuron’s responses was explained by the auditory stimulus (Fig 3A). For each responsive neuron, we computed the distribution of its sound-evoked responses across trials and then quantified how much auditory information it encoded by computing the mutual information between responses and stimuli *I*(*resp*; *stim*) [37–40]. Mutual information approaches zero when a neuron’s activity is independent of sound frequency and intensity, and increases as responses become more stimulus-driven. To compare the influence of BN on information across conditions and subpopulations, we calculated an information modulation index (IMI) ranging from -1 to 1, which indicates whether a neuron conveyed more information about the stimuli in No-BN or BN (Fig 3A). IMIs near 1 reflect greater encoding in BN, whereas IMIs near -1 indicate a loss of stimulus information in BN (Fig 3B). All subpopulations contained neurons with positive and negative IMIs, revealing diverse BN-induced changes in single-neuron encoding (Fig 3C, Kruskal-Wallis test, main effect for sub-population: *p* = 6.5648 × 10^−12^). On average, L2/3 neurons exhibited more negative IMIs than L5 IT and ET populations (Fig 3D; Dunn–Šidák post hoc test; L2/3-L5 IT *p* = 3.5019 × 10^−12^; L2/3-L5 ET *p* = 0.0011), whereas the two L5 subpopulations did not differ from each other (Dunn-Šidák post hoc test, L5 IT-L5 ET *p* = 0.1425). These findings indicate that L2/3 neurons reduce stimulus-related information in BN, whereas L5 IT and ET neurons maintain relatively preserved, more noise-invariant, response distribution. Together, these results show that BN induces stronger response suppression and reduced information encoding in L2/3, while L5 IT and ET neurons preserve balanced tuning and noise-invariant representations.

**Figure 3:**
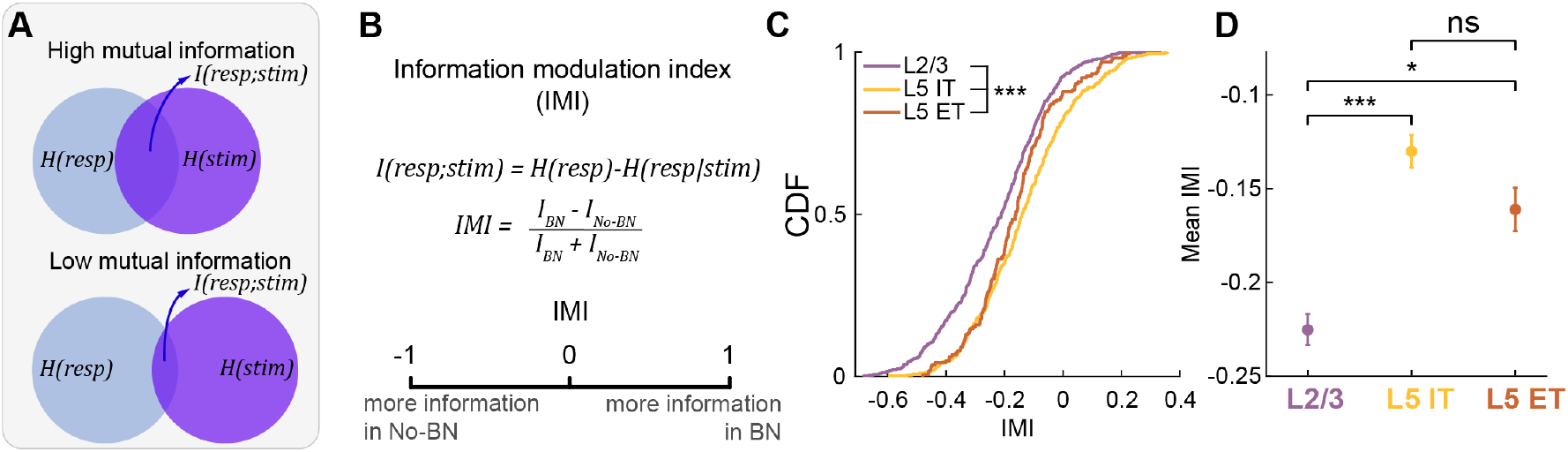
L2/3 single neurons encode less stimulus information in BN. **(A)** Schematic illustrating mutual information between neural responses and the auditory stimulus, *I*(*resp*; *stim*), in relation to the entropy of each variable. **(B)** Mathematical formulation of mutual information between neural activity and stimulus identity, along with the definition of the information modulation index (IMI). **(C)** Cumulative distributions of IMI values for each excitatory subpopulation (L2/3: *n* = 423, L5 IT: *n* = 290, L5 ET: *n* = 163). **(D)** Mean IMI values corresponding to the distributions in **C**. Same *n* as in **C**. Error bars denote mean ± s.e.m.

### BN reduces shared neural variability across spatial scales in IT but not ET neural responses

While single-neuron analyses reveal how individual responses change with BN, they do not capture how activity is coordinated within the densely interconnected cortical circuit. To determine whether BN alters coordinated activity between neurons, we quantified pairwise functional connectivity using two complementary measures: noise correlations and signal correlations.

If BN alters local circuit interactions, its effects should be reflected in the spatial organization of pairwise noise and signal correlations. Noise correlations quantify the shared trial-to-trial variability between two neurons after removing stimulus-driven activity, providing an estimate of their functional coupling [41–48]. To test whether BN modulates this coupling, we examined how noise correlations varied as a function of intersomatic distance (see Methods), to test whether BN preferentially affects functional coupling in local subnetworks or broadly across ACtx.

Consistent with previous studies [47, 49], the strongest noise correlations were observed at short intersomatic distances (Fig 4C, two-way ANOVA, main effect for intersomatic distance, L2/3: *p* = 1.5101×10^−51^, L5 IT: *p* = 7.5697×10^−28^, L5 ET: *p* = 2.3240×10^−11^). BN reduced noise correlations in L2/3 and L5 IT neurons, but not in ET neurons (Fig 4C, two-way ANOVA, main effect for BN, L2/3: *p* = 0.0089, L5 IT: *p* = 0.0022, L5 ET: *p* = 0.1302). In all subpopulation, this BN-induced reduction did not depend grossly on intersomatic distance, as indicated by non-significant interactions (Fig 4C, two-way ANOVA, interaction between BN and intersomatic distance, L2/3: *p* = 0.9398, L5 IT: *p* = 0.9768, L5 ET: 0.4861).

**Figure 4:**
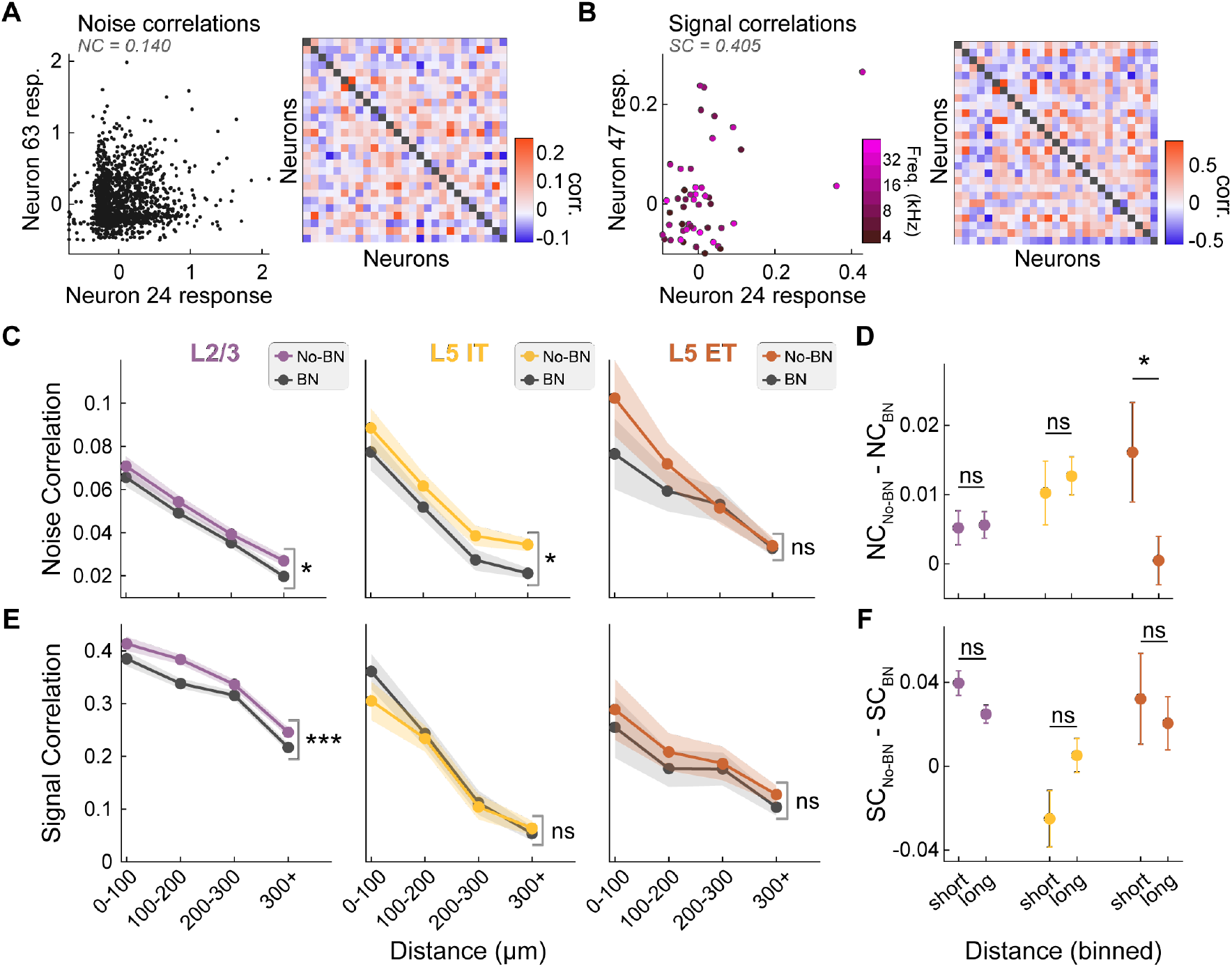
Excitatory subpopulations exhibit BN-dependent changes in pairwise correlations. **(A)** Mean-subtracted trial-by-trial responses for an example pair of simultaneously recorded neurons (left) and the corresponding noise correlation matrix for all neuron pairs within an example field of view (FoV; right). **(B)** Same as **A**, but showing signal correlations computed from tuning curves for the same example neurons. **(C)** Mean noise correlations plotted as a function of intersomatic distance under No-BN and BN conditions (L2/3: *n* = 9266, L5 IT: *n* = 3720, L5 ET: *n* = 2258 neuron pairs). **(D)** BN-induced change in noise correlations at short (≤200 *µ*m) and long (*>*200 *µ*m) intersomatic distances. **(E)** Same as **C**, but for signal correlations (same *n* as in C). **(F)** Same as **D**, but for signal correlations (same *n* as in D). Shaded regions denote mean ± s.e.m.

However, because the BN-induced reduction in noise correlations was asymmetric across distance, with larger effects at short distances and smaller effects at long distances (Fig 4C, right), we assessed whether BN differentially affected noise correlations across spatial scales by comparing BN-induced changes at short (≤ 200*µm*) and long distances (*>* 200*µm*). L5 ET neurons showed a stronger BN-induced reduction in noise correlations at short distances (Fig 4D, Wilcoxon rank sum test, L5 ET: *p* = 0.0285), whereas L2/3 and L5 IT neurons showed similar reductions at short and long distances (Fig 4D, Wilcoxon rank sum test, L2/3: *p* = 0.9239, L5 IT: *p* = 0.8463). These results indicate that BN broadly reduces shared variability across spatial scales in L2/3 and L5 IT populations, but acts more locally within L5 ET networks.

Next, we assessed pairwise functional connectivity through signal correlations, defined as the correlation between the tuning curves of two neurons (Fig 4B). BN induced a decrease in signal correlations only in L2/3 neurons (Fig 4E left, two-way ANOVA, main effect for BN, L2/3: *p* = 9.3076 × 10^−6^), whereas L5 IT and L5 ET neurons showed no significant change between BN conditions (Fig 4E center and right, two-way ANOVA, main effect for BN, L5 IT: *p* = 0.3666, L5 ET: *p* = 0.3435). For all subpopulations, the effect of BN on signal correlations did not differ between short and long intersomatic distances (Fig 4F, Wilcoxon rank-sum test, L2/3: *p* = 0.1262, L5 IT: *p* = 0.1289, L5 ET: *p* = 0.4635). Taken together with our prior analyses, these results indicate that BN not only reduces L2/3 response amplitudes, but also makes their tuning curves less similar to one another, whereas L5 IT and L5 ET neurons maintain stable tuning similarity across BN conditions.

### Population-level decoding of pure tones are noise-invariant in L5 IT

Although single-neuron activity and pairwise correlations reveal how BN modulates inputs to individual neurons and pairs, neural representations ultimately arise from the collective output of entire populations [50–54]. We therefore asked two key questions regarding how BN influences population-level stimulus representations: 1) does BN reduce the ability of neural populations to detect the presence of a pure tone, and 2) does BN impair their ability to discriminate between different pure tone frequencies?

To evaluate the noise invariance of population-level representations, we performed two decoding analyses to address both detection and discrimination of pure tones. We trained classifiers on trialwise population activity to test whether each subpopulation could reliably detect the presence of a sound and identify its frequency across BN conditions. To capture potential nonlinear interactions among neurons and to maintain a consistent architecture across decoding analyses, we first trained artificial neural networks to classify whether a sound was present on each trial based solely on sound-evoked activity (see Methods) (Fig 5A). Because classification performance depends on the number of neurons provided to the classifier, we trained all classifiers on randomly selected subsets of 40 neurons. This choice balanced the need for sufficient neurons to support reliable classification while avoiding oversampling in smaller FoVs, ensuring robust and comparable performance across FoVs. Each network consisted of an input layer of 40 units, two hidden layers of 16 units, and a single readout unit. Hidden layers used a *ReLU* activation function, and the readout was passed through a sigmoid nonlinearity, with outputs above 0.5 classified as sound-present. We cross-validated all decoding analyses by separating trials into independent training and testing sets and trained independent classifiers for each tone frequency. This design ensured that any reduction in classification performance under BN reflected diminished sound detection, rather than reductions arising from mismatches in frequency tuning between the trials used in the training and testing sets. For some example FoVs, detection performance in BN was markedly reduced (Fig 5B). Comparing average detection performance between No-BN and BN sessions revealed that L2/3 and L5 ET neurons exhibited reduced detection performance under BN, whereas L5 IT neurons showed noise-invariant detection of pure tones (Fig 5D, paired t-test, L2/3: *p* = 6.8345 × 10^−7^, L5 IT: *p* = 0.4586, L5 ET: *p* = 0.0073).

**Figure 5:**
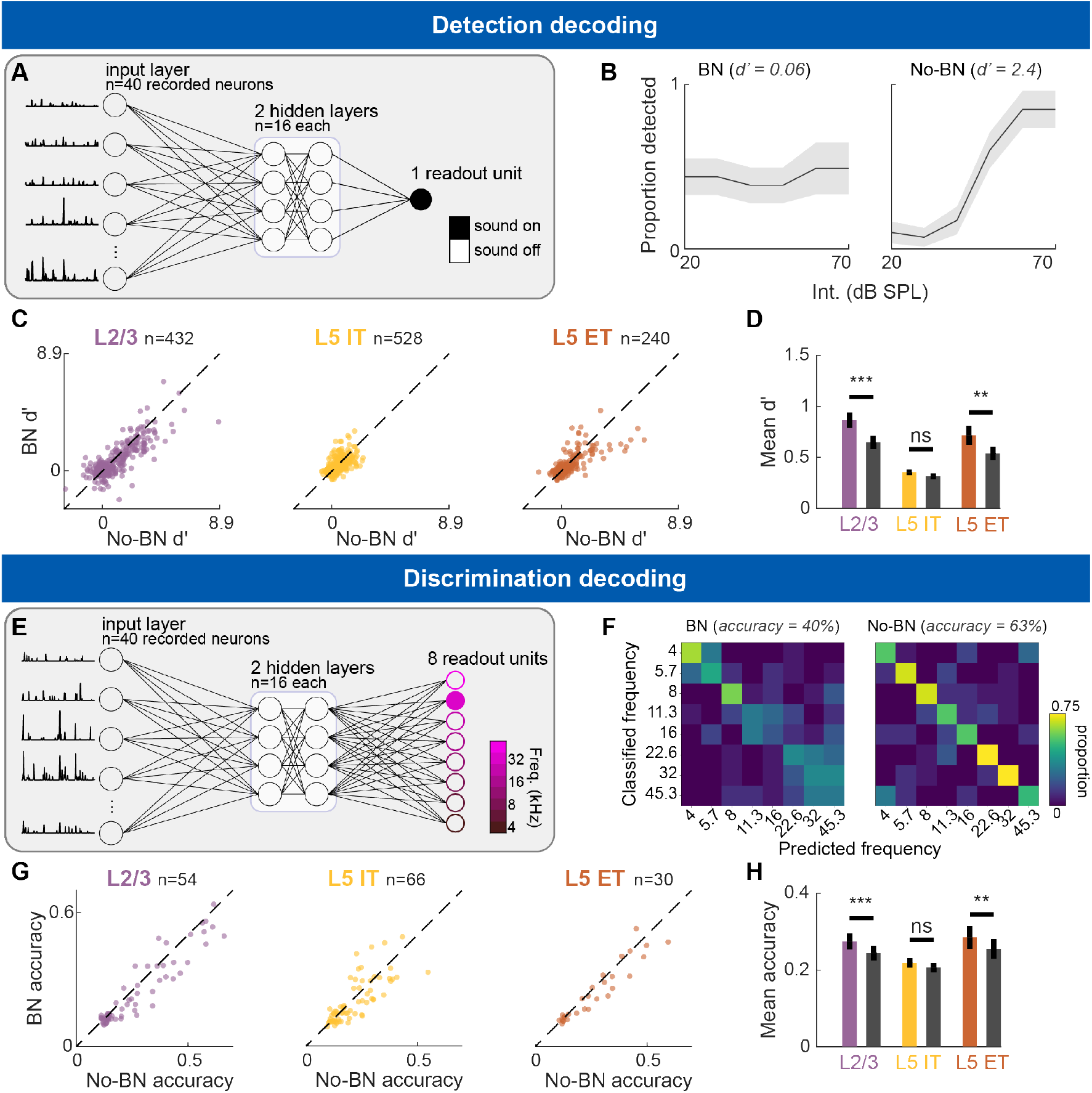
L5 IT neurons exhibit noise-invariant detection and discrimination of auditory stimuli. **(A)** Schematic of the artificial neural network used for binary detection decoding. **(B)** Example neurometric curves for the same neural population under BN (left) and No-BN (right) conditions. **(C)** Paired scatter plot of cross-validated detection performance (d’) for each frequency–intensity combination under No-BN and BN conditions. *n* denotes the number of binary classifiers. **(D)** Mean detection performance (d’) for each excitatory subpopulation, averaged across decoding runs shown in **C. (E)** Schematic of the artificial neural network used for multinomial discrimination decoding. **(F)** Example confusion matrices for the same neural population under BN (left) and No-BN (right) conditions. **(G)** Paired scatter plot of cross-validated discrimination accuracy for each stimulus intensity under No-BN and BN conditions. *n* denotes the number of multinomal classifiers. **(H)** Mean discrimination accuracy for each excitatory subpopulation, averaged across decoding runs shown in **G**. Error bars denote mean ± s.e.m.

To answer whether ACtx neural subpopulations can reliably discriminate pure tone frequencies, we next trained neural networks to classify each trial into one of the eight possible frequencies (Fig 5E). To make discrimination decoding directly comparable to detection decoding and to allow for the same degree of nonlinear interaction in the hidden layers, we used an identical architecture except for the readout: the output layer contained eight units instead of one. Each output unit corresponded to a single frequency, and a softmax function was applied to normalize the outputs and select the decoded frequency on each trial (Fig 5F). In this multinomial task, decoding accuracy remained stable for L5 IT neurons but decreased significantly for L2/3 and L5 ET neurons (Fig 5H, paired t-test, L2/3: *p* = 2.0953 × 10^−4^, L5 IT: *p* = 0.1521, L5 ET: *p* = 0.0026). Thus, L5 IT neural populations preserve both detection and discrimination decoding performance across BN conditions, while both L2/3 and L5 ET neural populations show reduced decoding performance for both sound detection and frequency discrimination in BN.

### L5 IT neurons maintain neural manifold geometry

While population-level decoding provides an estimate of how well each subpopulation encodes the specific pure-tone features used here, it does not address how many additional features (for example, more frequencies or stimulus dimensions) the same population could, in principle, encode. To assess the impact of BN on the structure and capacity of neural representations, we examined the geometry of population activity across conditions. Manifold geometry analysis [55] provides a way to quantify the efficiency of a population code by estimating the capacity of a neural population to represent multiple perceptual features.

Recent work has shown that neural population activity can be described in terms of neural manifolds and their effective dimensionality, which capture the dominant structure of population responses [55–57]. When sound-evoked responses are embedded in an *n*-dimensional space (one axis per neuron), activity trajectories typically lie on a lower-dimensional manifold rather than filling the entire space [50, 58]. Within this framework, responses to different stimuli (e.g., pure tone frequencies) form distinct manifold objects in neural space, each consisting of the set of response patterns associated with a given stimulus. The geometry of these manifold objects, particularly their size and dimensionality, constrains how efficiently a population can represent multiple stimuli [55]. For example, in a simulated population of three neurons, responses to three different stimuli might form manifold objects that resemble a line, disk, or sphere (Fig 6A, top). Lower-dimensional objects (e.g., lines and disks embedded in three dimensions) can be packed more efficiently in neural space, which promotes separability between manifold objects and increases the number of objects that can be encoded simultaneously. In contrast, larger or higher-dimensional objects quickly constrain the population representational capacity by making manifold objects less linearly separable [55] (Fig 6A, bottom). Thus, smaller and lower-dimensional manifolds correspond to a more efficient neural code, while larger and higher-dimensional manifolds reflect poorer encoding.

**Figure 6:**
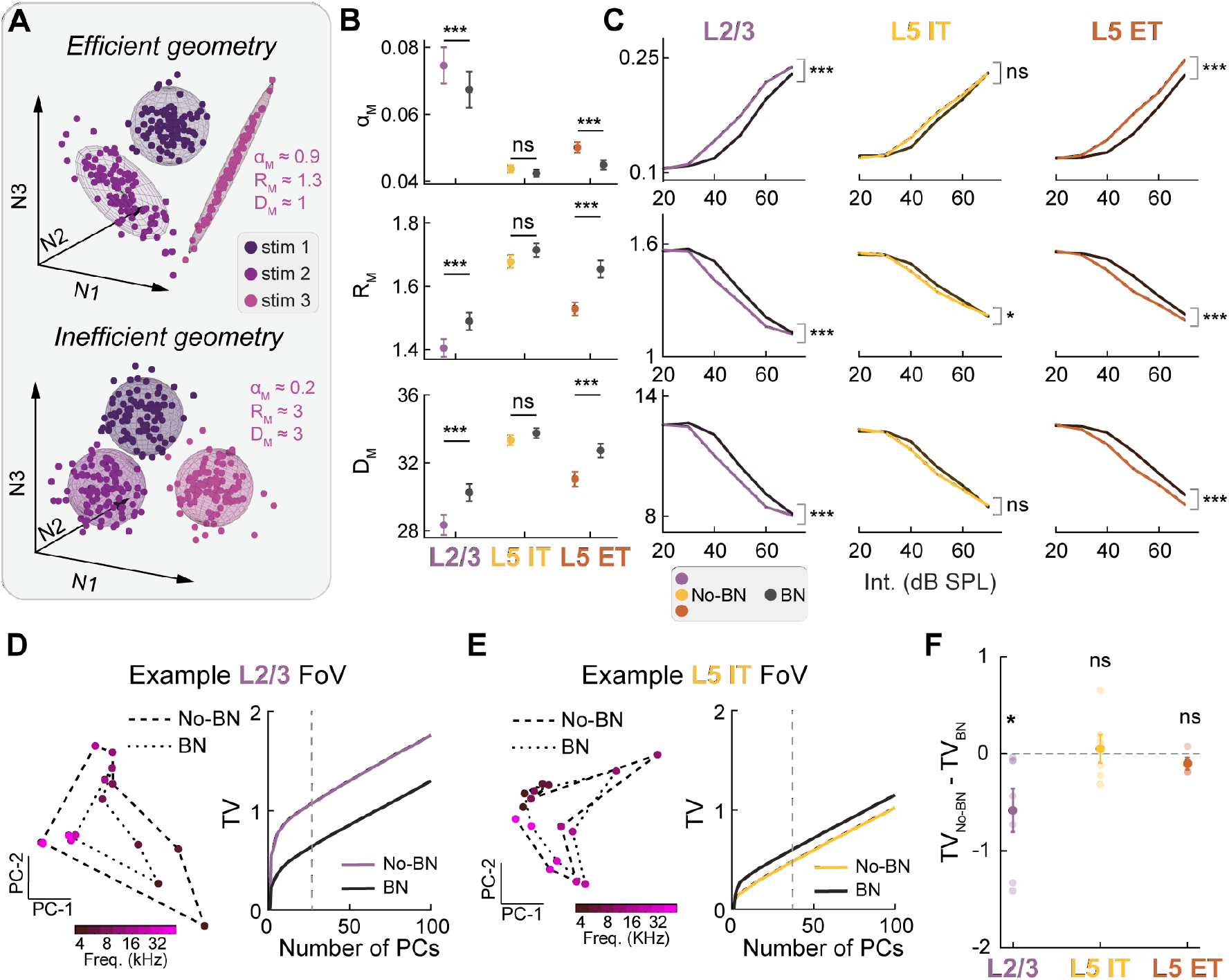
L5 ET neurons show noise-dependent manifold geometry but noise-invariant manifold size. **(A)** Simulated examples of manifold objects encoded by three neurons, illustrating differences in manifold geometry. Insets show the manifold capacity (*α*_*M*_), radius (*R*_*M*_), and dimensionality (*D*_*M*_) for the highlighted light pink manifold object in both examples. **(B)** Manifold geometry metrics for each excitatory subpopulation, with manifold objects constructed by pooling all intensities for each frequency (L2/3: *n* = 961, L5 IT: *n* = 2576, L5 ET: *n* = 566 neurons). **(C)** Manifold geometry metrics as a function of stimulus intensity, with manifold objects constructed from individual frequency–intensity combinations. Same *n* as in **B. (D)** Example sound-evoked population activity from a L2/3 FoV projected onto the first two principal components (left), and total variation of the population activity across principal components for each BN condition (right). The dotted line indicates the number of principal components required to explain 60% of the variance for this FoV. **(E)** Same as **D**, shown for an example L5 IT FoV. **(F)** Difference in total variation between No-BN and BN conditions for each FoV, shown separately for each subpopulation (L2/3: *n* = 10, L5 IT: *n* = 11, L5 ET: *n* = 5 FoVs). Error bars denote mean ± s.e.m.

Using this framework, we tested how BN affects manifold geometry across excitatory subpopulations. We quantified three established metrics of manifold structure: capacity (*α*_*M*_), radius (*R*_*M*_) and dimensionality (*D*_*M*_) which together reflect the efficiency of population-level encoding [55, 56, 59]. Effective encoding is characterized by high *α*_*M*_, low *R*_*M*_, and low *D*_*M*_, whereas reduced efficiency is marked by lower capacity and larger, higher dimensional manifolds (Fig 6A). In this simplified example, the neural manifold is composed of three neurons, which sets the maximum possible manifold dimensionality *D*_*M*_ to 3.

We first constructed manifold objects by pooling responses across all intensities for each frequency under No-BN and BN conditions, yielding eight objects per condition. Because manifold metrics depend on the dimensionality of the neural population, we analyzed randomly selected subsets of 40 simultaneously recorded neurons and resampled across subsets to approximate trends in the full population, which sets the maximum manifold dimensionality *D*_*M*_ to 40. L2/3 and L5 ET neurons showed significant BN-dependent changes in all three metrics (Fig 6B, Wilcoxon sign-rank test for *α*_*M*_, *R*_*M*_ and *D*_*M*_, respectively, L2/3: *p* = 4.2940 × 10^−7^, *p* = 4.3582 × 10^−8^, *p* = 3.3680 × 10^−9^, L5 ET: *p* = 6.8053 × 10^−10^, *p* = 6.462 × 10^−10^, *p* = 2.6811 × 10^−10^). In contrast, L5 IT neurons showed no significant BN-related differences in manifold capacity, radius, or dimensionality (Fig 6B, Wilcoxon sign-rank test for *α*_*M*_, *R*_*M*_ and *D*_*M*_, L5 IT: *p* = 0.1148, *p* = 0.1488, *p* = 0.1363).

To examine the effects of BN on manifold geometry at a finer stimulus scale, we repeated this analysis using one manifold per frequency–intensity combination, yielding 48 manifold objects per BN condition. This allowed us to assess how BN affected the geometry of representations for specific acoustic stimuli. L2/3 and L5 ET manifolds shifted toward less efficient geometry (Fig 6, two way ANOVA, main effect of BN for *α*_*M*_, *R*_*M*_ and *D*_*M*_, respectively, L2/3: *p* = 7.5707 × 10^5^, *p* = 4.2740 × 10^−6^, *p* = 4.5530 × 10^−7^, L5 ET: *p* = 6.7232 × 10^−5^, *p* = 2.1831 × 10^−7^, *p* = 7.7084 × 10^−8^). In contrast, L5 IT manifolds showed noise-invariant capacity and dimensionality, with BN selectively increasing manifold radius (two way ANOVA, main effect of BN for *α*_*M*_, *R*_*M*_ and *D*_*M*_, respectively, L5 IT: *p* = 0.1097, *p* = 0.0329, *p* = 0.0546). These results demonstrate that the manifold geometry of L2/3 and L5 ET populations is particularly susceptible to BN, while L5 IT neurons preserve a more noise-invariant manifold structure.

Although manifold geometry analysis characterizes the structure of individual manifold objects, it does not capture the global spread of all objects within the neural representational space. To assess whether BN alters this global dispersion of sound-evoked population activity, we first visualized the data using principal component analysis (PCA; see Methods). In some FoVs, the first two principal components (PCs) revealed a clear contraction of the span of stimulus representations under BN (Fig 6D, left), while in others the overall spread appeared unchanged (Fig 6E, left).

To quantify these differences, we identified for each FoV the minimum number of PCs that explained at least 60% of the variance in sound-evoked activity under both BN conditions (Fig 6D,E, right). We then projected the data into this reduced space and computed the manifold size as the total variation (TV) of the projected activity (see Methods). TV provides a robust scalar measure of the effective radius of the point cloud in lower-dimensional projections of varying dimensionality, enabling comparisons across FoVs. This analysis revealed a significant reduction in manifold size for L2/3 populations under BN (Fig 6F, Wilcoxon signed rank test, L2/3: *p* = 0.0020), while the L5 IT and ET populations did not show a change in TV under BN (Wilcoxon signed rank test, L5 IT: *p* = 0.5771, L5 ET: *p* = 0.1250). Thus, the global representational space contracted in L2/3 under BN, while remaining effectively noise invariant in L5 IT and L5 ET populations.

These two analyses, manifold geometry and manifold size, revealed an apparent discrepancy for L5 ET. In L2/3, BN both reduced the efficiency of manifold geometry and contracted overall manifold size, whereas L5 IT remained noise-invariant in both measures. In contrast, L5 ET neurons showed less efficient manifold geometry under BN, but preserved their global manifold size. Although this pattern is difficult to visualize directly, it implies that individual manifold objects became larger and more high-dimensional without increasing the overall spread of the full set of objects. This could occur if manifolds expand inward (toward the origin) or toward one another rather than outward, two possibilities that are not mutually exclusive. This scenario is consistent with the decoding results (Fig 5D,H), where L5 ET detection and discrimination performance decline under BN, as would be expected if near-boundary trials are displaced toward neighboring frequencies or toward the origin in neural space. Overall, BN degrades the efficiency of population-level encoding within manifold objects in L5 ET, while the total neural subspace used for population-level representations remains unchanged.

## Discussion

We imaged sound-evoked responses in defined excitatory subpopulations of the ACtx with *in vivo* two-photon microscopy. To distinguish how different circuit elements contribute to noise invariance, we compared an excitatory subpopulation primarily involved in local processing (L2/3) with two deep-layer subpopulations that broadcast information to distant targets (L5 IT and L5 ET). All three subpopulations adjusted their responses when tones were embedded in BN, but noise invariance was concentrated in the broadcast pathways. L2/3 neurons showed clear noise dependence, including suppressed single-neuron responses (Fig 2, Fig 3), increased pairwise correlations (Fig 4), and reduced fidelity in population-level encoding of tone identity (Fig 4, Fig 5, Fig 6). Under identical experimental conditions, L5 IT neurons maintained stable single-neuron and population-level representations, with noise-related differences appearing only in pairwise correlations (Fig 4C). L5 ET neurons expressed a more limited form of noise invariance: their single-neuron and pairwise responses were largely stable across BN conditions, whereas their population decoding performance and manifold structure were not. Together, these results show that excitatory subpopulations in ACtx rely on different representational levels to preserve sensory information in noise and reveal a functional stratification of noise invariance across the cortical microcircuit.

### Significance of noise invariance for sound processing

ACtx plays a critical role in extracting behaviorally relevant sound features [6, 22, 60–62], and the canonical cortical microcircuit carries and transforms sensory information within ACtx [28]. In this circuit, thalamocortical inputs primarily innervate L4, which relays information to L2/3, then to L5 and L6, where signals are broadcast widely throughout the brain [25, 26, 29, 63]. L2/3 neurons are the first major cortical recipients of L4 input and provide dense, complex projections to L5 [24], a principal output layer with extensive long-range targets [29, 64–69]. Along the ascending auditory pathway, sound representations become increasingly noise-invariant from the periphery to ACtx [1], and even more so in higher-order ACtx [19]. When considered together, the hierarchical organization from periphery to cortex and from L4 to L2/3 to L5 supports a model in which ACtx integrates noise-dependent inputs, refines them through subpopulation-specific computations, and broadcasts increasingly noise-invariant representations of sounds that can drive adaptive behavior [28, 70, 71].

Prior studies have examined sensory representations that remain invariant to factors other than BN. For example, work on level invariance shows that ACtx representations can remain stable despite changes in sound intensity [72, 73], and non-primary ACtx can form distractor-invariant representations of sounds when they are behaviorally relevant [74, 75]. In the visual system, a rich body of work has investigated how neurons in the visual pathway support object recognition independently of changes in rotation, position, and other variables [76–80]. A common limitation of many of these studies is that they treat cortical excitatory neurons as a homogeneous population, overlooking the heterogeneity of subpopulations within a cortical column. In light of our findings, this gap raises the possibility that invariant coding in other sensory modalities may also arise from specialized computation by distinct excitatory subpopulations.

In this study, we used pure tones as the signal and white noise as the masker. Pure tones give precise control of frequency and intensity to isolate mechanisms of noise invariance and is a common stimulus choice in work probing noise-invariant representations in the auditory system [2, 3, 6, 10, 81], but they sample only a small region of the space of natural sounds. Extending our findings to more complex stimuli will require careful stimulus design. For example, time-varying stimuli, such as amplitude-modulated tones, introduce fluctuations in SNR over time and would require ACtx neurons to express noise invariance both for spectral content and for temporal envelope tracking. Human studies suggest that envelope tracking can remain relatively preserved in BN [82], but the contribution of ACtx neurons to time-varying noise invariance is not yet known. Stimuli with richer spectral structure, such as frequency sweeps and natural vocalizations, would demand invariance across multiple spectral and temporal channels. Although ACtx neurons exhibit multiplexed responses to several sound features [38, 75, 83–85], future work should test whether individual neurons maintain a consistent degree of noise invariance across different regions of their receptive fields and whether population-level codes support multiplexed forms of noise invariance across features and timescales.

### Differences in noise invariance between L5 subpopulations

At the level of individual neurons, L5 IT and ET populations exhibited a similar degree of noise invariance in both tuning and response distributions (Fig 2B,G). However, differences become more pronounced in pairwise and population-level representations. Pairwise analyses showed that L5 IT neurons exhibited reduced noise correlations across all intersomatic distances, whereas L5 ET neurons showed reductions primarily only locally (Fig 3C-D). Noise correlations can reflect shared inputs, functional coupling, and the information content of population responses [86–88]. The broad reduction in L5 IT (and L2/3) correlations suggests more global changes in correlated variability under BN, whereas the spatially restricted effects in L5 ET may reflect BN-dependent modulation of local subnetworks within this subpopulation.

Population-level differences were even more striking. Decoding analyses showed that BN impaired both sound detection and frequency discrimination in L5 ET neurons, whereas L5 IT decoding performance remained stable (Fig 5D,H). Consistent with this result, manifold geometry analyses revealed that BN disrupted the fine-scale structure of sound representations in L5 ET neurons, altering manifold capacity, radius, and dimensionality, while affecting only manifold radius in L5 IT neurons (Fig 6C). Notably, despite these changes in fine-scale geometry, the global structure of sound representations in both L5 IT and L5 ET populations remained noise invariant, as reflected by stable manifold size across BN conditions (Fig 6F).

In L5 ET neurons, this dissociation between degraded fine-scale geometry and preserved global structure suggests that BN may cause individual manifold objects to expand toward one another or toward the origin, without changing the overall extent of the population-level representation. Together, these results indicate that L5 ET neurons preserve the global size of their neural manifold under BN but exhibit degraded stimulus-specific geometry, whereas L5 IT neurons maintain both global and fine-scale structure. This divergence may reflect differences in the functional demands of their downstream targets. If L5 IT and L5 ET projection targets differ in computational requirements or modularity, the corresponding cortical output pathways may differentially shape population responses to preserve noise-invariant representations appropriate for their target circuits.

These findings add to a growing body of evidence that L5 IT and L5 ET neurons are functionally distinct. Across cortical areas, these subpopulations differ in their projection targets as well as in multiple morphological and physiological properties [21, 28, 29, 65, 89, 90]. Here, we show that although both L5 IT and L5 ET neurons participate in broadcast pathways and exhibit noise invariance at multiple representational levels, they differ in the extent to which specific pairwise and population-level metrics remain noise invariant. One possible explanation for this differential modulation by BN is that L5 IT and L5 ET neurons receive distinct long-range inputs [29, 65]. Differences in top-down modulation could lead to subpopulation-specific effects of BN by selectively enhancing or suppressing neuronal responses, thereby shaping the degree of noise invariance expressed at the population level. An additional possibility is that these subpopulations differ in their local circuit organization, including recurrent connectivity within each group and their interactions with other neural populations, such as inhibitory interneurons [67, 91, 92]. Together, differences in long-range inputs, local recurrence, and inhibitory interactions may underlie the distinct patterns of noise invariance observed between L5 IT and L5 ET neurons. Future studies could directly test these ideas by transiently inactivating cortical regions that provide top-down input to L5 neurons, such as posterior parietal cortex or anterior cingulate cortex, and by using cell-type-specific optogenetic perturbations to assess whether local microcircuits are differentially engaged in the presence of BN.

### Potential mechanisms that lead to noise invariance

Early work on noise invariance emphasized mechanisms in the auditory periphery that reduce the overall gain of auditory nerve responses [11, 93]. Although such mechanisms can improve sound coding in noisy environments, noise-dependent distortions of sound representations remain evident beyond the periphery, indicating that global gain adjustments alone are insufficient to account for noise-invariant coding. Instead, peripheral adaptations likely constitute the initial stage of a multistep computation that is progressively refined in downstream auditory structures.

Cholinergic projections from the basal forebrain exhibit strong layer-, subpopulation-, and region-specific organization. Across cortical areas, basal forebrain inputs exert layer-specific effects [94, 95], and within ACtx, cholinergic innervation differs between primary and non-primary subdivisions [17, 95]. At the cellular level, cholinergic signaling differentially modulates L5 IT and L5 ET neurons [66, 96]. In parallel, sound representations become progressively more noise invariant across layers of the cortical microcircuit in primary ACtx and between primary and non-primary auditory fields, mirroring differences in cholinergic innervation and functional responses across these populations. Consistent with this framework, recent work has implicated cholinergic input to ACtx as a potential mechanism supporting noise-invariant representations, in part through its effects on spontaneous firing rates and local synchrony [3]. Together, these observations suggest that neuromodulatory influences may contribute to subpopulation-specific differences in noise invariance across the auditory hierarchy and within the ACtx microcircuit.

In addition to neuromodulatory influences, inhibitory circuits have been proposed as mechanisms contributing to the construction of noise-invariant representations in ACtx. Inhibitory neurons play well-established roles in sensory processing, including surround suppression [5, 97, 98], suppressive feedback [99, 100], and temporal sharpening [101, 102]. With respect to noise invariance, inactivation of parvalbumin (PV) or somatostatin (SOM) interneurons impairs behavioral performance in noisy conditions to a degree comparable to inactivation of ACtx itself [6]. This result indicates that PV and SOM activity is necessary for detecting sounds in BN, but also suggests that inhibitory neurons alone do not fully account for the cortical mechanisms underlying noise-invariant perception. Consistent with this interpretation, optogenetic activation of PV neurons suppresses ACtx tuning curves in a manner similar to BN; however, combining PV activation with BN produces even stronger suppression, indicating that PV activity alone is insufficient to explain the full modulation of tuning curves observed in noisy environments [2].

Notably, many of these studies have treated inhibitory neurons as a homogeneous population within ACtx. In contrast, both theoretical and experimental work suggests that feedforward inhibition may support contrast gain control within the canonical cortical microcircuit [71], potentially supporting noise invariance by selectively modulating the gain of individual neurons according to their receptive fields and bottom-up inputs. Under this framework, inhibitory neurons would themselves be differentially engaged by BN and would, in turn, selectively enhance or suppress excitatory neurons within the same layer, thereby stabilizing sound representations in noisy conditions. A key prediction of this mechanism is that noise invariance should increase across the cortical microcircuit, from L2/3 to L5, a pattern that is consistent with our findings. Future experiments could directly test this hypothesis by identifying interneuron populations whose activity is selectively modulated by BN and determining whether their trial-by-trial influence on local excitatory neurons adjusts gain in a manner that promotes noise-invariant sound representations.

Our results demonstrate that excitatory subpopulations in ACtx make distinct and complementary contributions to constructing noise-invariant representations. We speculate that a complete mechanism for noise invariance within ACtx must include components that differentially influence the three excitatory subpopulations examined here and can account for the mixed pattern of invariance across single-neuron, pairwise, and population levels. Such components may include finely tuned gain control by inhibitory interneurons, as well as neuromodulatory inputs that are selectively engaged in the presence of BN.

### Conclusions

Disentangling sensory signals from background noise is a fundamental process that enables animals to represent stimuli accurately and generate appropriate behavioral responses. We show that excitatory subpopulations in ACtx respond differentially to sounds in BN, depending on their laminar position and projection class within the cortical microcircuit. This subpopulation-specific organization supports the idea that deep-layer broadcast pathways preferentially carry noise-invariant representations, whereas superficial populations remain more noise-dependent. Our findings bridge the noise-dependent representations observed in earlier stages of the auditory pathway with the more noise-invariant representations reported in higher auditory areas. Together, they underscore the role of excitatory subpopulations in implementing the computations that give rise to noise-invariant coding.

## Materials and Methods

### Mice

All procedures were approved by the University of Pittsburgh Animal Care and Use Committee and follow the National Institute of Health guidelines for the care and use of laboratory animals. Data were collected from 17 mice (10-16 weeks old, both male and female). For L2/3 recordings, we used two C57BL/6 mice (#000664, Jackson Labs), and five Emx1-Cre mice (#005628, Jackson Labs). For L5 IT recordings, we used six Tlx3-Cre mice (B6.FVB(Cg)-Tg(Tlx3-Cre)PL56Gsat/Mmucd, MMRRC). For L5 ET mice, we used four C57BL/6 mice. All mice were housed on a 12 h light/dark cycle with ad libitum access to food and water. All imaging was conducted during the dark cycle.

### Surgical Procedures

#### Virus-mediated gene delivery

Mice were anesthetized with 4% isoflurane and positioned in a stereotaxic frame (model 1900, Kopf). Throughout the procedure, a surgical plane of anesthesia was maintained using a continuous infusion of isoflurane (2%) in oxygen. Mice lay atop a homeothermic blanket system (Fine Science Tools) that maintained core body temperature at approximately 36.5°C. The scalp was shaved and sterilized with alternating applications of iodine and ethanol, followed by subcutaneous injection of lidocaine hydrochloride (5 mg/ml) for local analgesia.

For ACtx injections, a ∼1 cm incision was made between the right eye and ear to expose the temporalis muscle, which was then retracted. Two burr holes (∼0.3 mm diameter each) were drilled along the right temporal ridge, spanning a region 1.5–2.5 mm rostral to the lambdoid suture. For inferior colliculus (IC) injections, a midline incision was made to expose bregma and lambda. The skull was leveled such that the vertical difference between bregma and lambda was less than 100 *µ*m, and a single burr hole was drilled at 4.8 mm caudal and 0.9 mm lateral to bregma.

Viral injections were performed using a motorized stereotaxic injector (Nanoject III, Drummond Scientific). For ACtx injections, 250 nl of either a non-conditional GCaMP8s virus (pGP-AAV-syn-jGCaMP8s-WPRE, Addgene, titer: 3.5 x 10^12^ vg/mL) or a Cre-dependent GCaMP8s virus (pGP-AAV-syn-FLEX-jGCaMP8s-WPRE, Addgene, titer: 6 x 10^12^ vg/mL) was delivered at a depth of approximately 450 *µ*m below the pial surface at each injection site. For IC injections, 250 nl of retrograde GCaMP8s virus (pGP-AAV-syn-jGCaMP8s-WPRE, Addgene, titer: 4 x 10^12^ vg/mL) was delivered at depths of 900 *µ*m and 400 *µ*m below the pial surface. Following injections, the surgical sites were closed, antibiotic ointment was applied, and postoperative analgesia was administered subcutaneously (carprofen, 5 mg/ml). Mice were provided with ad libitum access to a carprofen MediGel and were closely monitored for three days following surgery.

#### Cranial window implantation

Mice were brought to a surgical plane of anesthesia using the same anesthesia and temperature-control procedures described above. The dorsal surface of the skull was exposed, and the periosteum was removed. The skull was cleaned with 70% ethanol and chemically etched before affixing a custom titanium head plate (eMachineShop). The head plate was secured to the skull with opaque dental cement (C&B Metabond) and allowed to fully cure. After head-plate attachment, the temporalis muscle was retracted to expose the temporal ridge. A circular outline (3 mm diameter) centered over the temporal ridge approximately 0.5 mm above the lambdoid suture was marked using a biopsy punch. The skull within and around this outline was thinned using a hand drill to create a flat surface. Once sufficiently thinned, the outlined bone was carefully removed with a scalpel to expose the underlying cortex. A cranial window was constructed by placing a stack of glass coverslips (two 3 mm diameter and one 4 mm diameter) over the exposed brain. The edges of the craniotomy were sealed with silicone elastomer (Kwik-Sil) to create an airtight seal, and the window was secured with opaque dental cement applied around the perimeter of the 4 mm coverslip. All remaining exposed skull was covered with dental cement, and the surrounding skin was affixed to the cement using Vetbond (3M) tissue adhesive. Mice recovered under the same postoperative analgesia and monitoring conditions used following viral injections.

### Acoustic Stimulation

Stimuli were generated with a 24-bit digital-to-analog converter (National Instruments model PXI-4461) using custom scripts written in MATLAB (MathWorks) and LabVIEW (National Instruments). Acoustic stimuli were delivered via a free-field speaker (PUI Audio) facing the left ear and calibrated using a free-field prepolarized microphone (377C01, PCB Piezotronics).

### Calcium Imaging

Light-reversed mice were awake and head-fixed for all recording sessions. Prior to imaging, mice were habituated to head-fixation and the recording chamber for several days. Neural activity in response to four pure tones (4, 8, 16, and 32 kHz) were captured by widefield fluorescence imaging (Bergamo, ThorLabs) and used to functionally confirm the location of the right primary ACtx. Two-photon calcium imaging was conducted using an InSightX3 (Spectra Physics) Laser tuned to 940 nm and a water-immersion objective (Nikon 16x). All two-photon imaging (Bergamo, ThorLabs) was of the right ACtx. Mice were head-fixed upright with the microscope rotated to be parallel to the cranial window (approximately 40 to 50^*◦*^ tilt). Images were collected at 30 Hz. The depth below pial surface used for recordings depended on neuron subtype (L2/3: 150-250 *µ*m, L5 IT: 350-500 *µ*m, L5 ET: 450-600 *µ*m). Separate FoVs from the same mouse were at least 50 *µ*m above or below the original imaging plane. All two-photon calcium imaging was conducted within a dark, sound-attenuating chamber.

Two imaging sessions were performed for each FoV: one without background noise (No-BN) and one with background noise (BN). Each session consisted of 960 trials, comprising 20 repetitions of 50 ms pure tones presented in pseudo-random order. Tones varied in frequency (4–45 kHz) and intensity (20–70 dB SPL), yielding 48 unique frequency–intensity combinations. Stimulation parameters were identical across sessions, with the sole difference being the presence of continuous white noise at 50 dB SPL delivered from a secondary speaker positioned directly below the primary stimulation speaker during BN sessions. Each mouse underwent as many imaging sessions as FoVs available. Each session lasted approximately 48 minutes, corresponding to a total of ∼88,000 imaging frames per session.

### Data Analysis

#### Image processing

Two-photon imaging data were processed using the open-source software Suite2P [34]. Image stacks were motion-corrected by rigid registration, and regions of interest (ROIs) were automatically detected with neuropil subtraction. All ROIs were manually curated to ensure that they corresponded to individual neurons. Calcium fluorescence traces were deconvolved to estimate spike rates and then z-scored within each session by subtracting the baseline mean and dividing by the baseline standard deviation. Imaging sessions from the same field of view were aligned between conditions using the ROICaT cell-matching algorithm. Suite2P outputs from paired sessions were registered to each other, and ROIs with overlapping spatial footprints were identified as matching neurons across sessions.

#### Responsiveness

Neuronal responsiveness was determined using an approach adapted from Kato et al. [103]. A neuron was considered responsive to a given stimulus if it met two criteria: (1) sound-evoked responses exceeded 0.5 z-scores above baseline in at least 50% of trials, and (2) the mean sound-evoked response across all trials exceeded 1 z-score. Responsiveness was assessed separately for each unique frequency–intensity combination and aggregated between both BN conditions. A neuron was classified as sound responsive if it met these criteria for at least one stimulus.

#### Tuning curves

Each recorded neuron was tested with 20 trials of each unique stimulus, yielding a total of 960 trials per session (48 frequency–intensity combinations). Neural activity within a fixed response window (0.5 s following sound onset) was averaged across trials to construct a frequency response area (FRA) for each neuron. FRAs were represented as matrices with eight rows corresponding to frequencies and six columns corresponding to intensities (Fig 2A). Frequency tuning curves were derived by averaging responses across all intensities for each frequency, and intensity tuning curves were derived by averaging responses across all frequencies for each intensity. To construct population-averaged frequency tuning curves, individual neuron tuning curves were centered on their best frequency (BF) and then averaged within each subpopulation. Intensity tuning curves were averaged across neurons without re-centering.

#### RMA regression

To compare tuning curves of individual neurons across BN conditions, we used reduced major axis (RMA) regression [18, 36]. Unlike ordinary least-squares regression, RMA accounts for measurement noise in both variables. For each neuron, RMA was applied separately to frequency and intensity tuning curves to estimate slope and intercept parameters. Only slope and intercept estimates that differed significantly from their null values (slope = 1, intercept = 0) were included in subsequent analyses. Statistical significance was assessed using one-sample t-tests comparing the estimated coefficients to their respective null values, with standard errors derived from the residual variance of the fitted model [104]. Intercepts significantly greater than or less than zero were classified as additive or subtractive shifts, respectively, whereas slopes significantly greater than or less than one were classified as multiplicative or divisive scaling.

#### Mutual information

Before computing mutual information, we empirically estimated the marginal and joint probability distributions *P* (*X*), *P* (*S*), and *P* (*X, S*). Neural responses *X* were discretized by binning sound-evoked activity using a fixed bin width of 0.2 z-scored responses. Stimulus identity *S* was encoded as an integer index ranging from 1 to the total number of unique stimuli (48) and had a uniform distribution by experimental design. The joint distribution *P* (*X, S*) was obtained by histogramming sound-evoked responses separately for each stimulus and normalizing across all trials.

Mutual information between the neural response and stimulus identity was computed as:

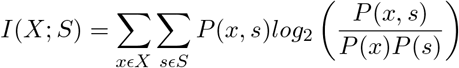

This calculation yielded a single non-negative scalar value quantifying how much information a neuron’s responses conveyed about stimulus identity. Mutual information was computed separately for the BN and No-BN conditions. Because the stimulus set represented only a limited sampling of the auditory space encoded by ACtx neurons, we applied a bias correction to each mutual information estimate. For each neuron, bias was estimated using the analytically derived approximation from Panzeri et al. [105]:

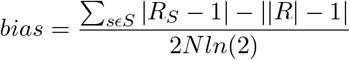

where |*R*| denotes the number of response bins with nonzero probability in *P* (*X*), |*R*_*s*_| denotes the number of response bins with nonzero probability in the conditional distribution *P* (*X* | *S* = *s*), and *N* is the total number of stimulus presentations. The estimated bias was subtracted from each mutual information value.

To quantify how background noise altered stimulus-related information on a neuron-by-neuron basis, we computed an information modulation index (IMI):

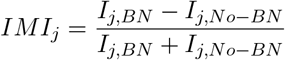

Each neuron was thus assigned an IMI value ranging from −1 to +1. Positive IMI values indicate greater stimulus-related information in BN than in No-BN, values near zero indicate little difference between conditions, and negative values indicate greater information encoding in No-BN.

#### Pairwise correlations

Noise correlations (spike-count correlations) were computed by subtracting, for each neuron, its mean response across trials for a given stimulus and then calculating the Pearson correlation coefficient between the resulting trial-by-trial residuals for each pair of simultaneously recorded neurons. Signal correlations (tuning correlations) were computed as the Pearson correlation between the corresponding entries of the FRAs of pairs of simultaneously recorded neurons. All correlation analyses were restricted to neurons classified as sound responsive.

#### Detection and discrimination decoding

We evaluated stimulus-encoded information at the population level using two decoding approaches: discrimination and detection. For detection decoding, we constructed training datasets consisting of sound-on and sound-off trials. Sound-on trials were drawn from responses to tones presented at 60 and 70 dB SPL for a single frequency, whereas sound-off trials consisted of randomly sampled neural activity measured 2 s after sound onset, when no stimulus was present. Decoders were trained separately for each frequency to avoid confounds arising from frequency tuning differences between training and testing data. We trained a neural network with the same architecture used for frequency discrimination decoding, except that the output layer consisted of a single artificial neuron. The output neuron was normalized to values between 0 and 1 using a sigmoid activation function, and trials were classified as sound-present if the output exceeded a threshold of 0.5.

Detection performance was quantified using d’ (d-prime), which incorporates both hit rate (*HR*) and false alarm rate (*FAR*):

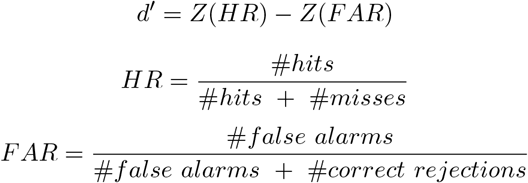

For discrimination decoding, we trained a neural network to classify stimulus frequency based on the activity of 40 simultaneously recorded neurons. Neural activity was passed through two hidden layers of 16 artificial neurons each and then into an output layer of eight artificial neurons, one corresponding to each stimulus frequency. Hidden layers used rectified linear unit (ReLU) activation functions, and the output layer was normalized using a softmax function. Network performance was assessed using ten-fold cross-validation, and decoding accuracy was quantified as the mean classification accuracy across folds. To control for intensity-dependent effects, discrimination decoding was performed using trials from a single sound intensity at a time and repeated independently for each intensity.

#### Manifold geometry

To characterize the population-level geometry of sound-evoked neural activity, we applied a mean-field-theoretic manifold geometry analysis [55]. This framework quantifies the classification capacity of a neural population by describing the geometric structure of population responses to different stimulus categories. For each frequency, we extracted population activity from a single FoV and represented each trial as a vector in R^*n*^, where each dimension corresponded to the activity of one neuron. To standardize dimensionality across FoVs and subpopulations, we analyzed randomly sampled subsets of *n* = 40 neurons and resampled with replacement to approximate the full recorded population. Trials corresponding to the same stimulus category formed a point cloud in neural space, which was treated as a single neural manifold. For each manifold, we extracted three geometric metrics: manifold radius (*R*_*M*_), effective dimensionality (*D*_*M*_), and classification capacity (*α*_*M*_). We performed this analysis in two complementary ways. First, we pooled trials across all intensities for each frequency, yielding one manifold per frequency (Fig 6B). Second, we constructed separate manifolds for each frequency-intensity combination, yielding a finer-grained analysis of population geometry (Fig 6C).

Manifold geometry analysis quantifies how efficiently population responses to different stimuli can be separated in high-dimensional neural space. The manifold radius *R*_*M*_ measures the spatial extent of trial-to-trial variability within a stimulus category, whereas the effective dimensionality *D*_*M*_ reflects the number of dimensions required to capture this variability. Together, these metrics describe how compact and low-dimensional a stimulus representation is, properties that facilitate separability from other stimulus manifolds. For example, if responses to a given frequency become more consistent across trials, the corresponding point cloud contracts toward a single point, resulting in smaller values of *R*_*M*_ and *D*_*M*_.

The manifold capacity *α*_*M*_ provides an integrated measure of coding efficiency by quantifying how many such manifolds can be linearly separated by the same neural population. In the mean-field framework, *α*_*M*_ is inversely related to both *R*_*M*_ and *D*_*M*_; thus, reductions in capacity indicate increased manifold size, increased dimensionality, or both. For our purposes, *α*_*M*_ serves as a compact summary metric linking population geometry to the efficiency of stimulus encoding.

#### Manifold size

To quantify the global size of the neural manifold, we first projected population activity into a lower-dimensional subspace that captured a substantial fraction of the variance in neural responses. For each neuron, we computed the peri-stimulus time–averaged activity for each unique stimulus (48 total), pooling trials from both No-BN and BN sessions. The activity of all simultaneously recorded neurons was then aggregated into a *t × n* matrix, where *t* denotes time points and *n* denotes the number of neurons. We reduced the dimensionality of this matrix using principal component analysis (PCA), projecting the data into a *t × p* subspace, where *p* was chosen as the minimum number of principal components required to explain at least 60% of the total variance. After projecting the neural activity into this reduced space, we quantified the overall size of the population representation by computing its total variation (TV), defined as

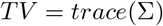

where Σ is the covariance matrix of the projected neural activity. TV provides a scalar measure of the overall spread of population responses in the reduced neural space and thus serves as an estimate of global manifold size.

### Statistical Analysis

All statistical analyses were performed in MATLAB (MathWorks). Data are reported as mean *±* s.e.m. unless otherwise noted. When data did not meet assumptions of normality, nonparametric statistical tests were used. For multi-group comparisons, post hoc tests were performed using Tukey’s or Dunn’s procedures, as appropriate. Statistical significance in figures is denoted as * *p <* 0.05, ** *p <* 0.01, and *** *p <* 0.0001.

## Acknowledgments

We thank current and former members of the Williamson Lab for helpful feedback and discussions. This work was supported by NIH/NIDCD grants R21DC018327 and R01DC020459, and the Klingenstein-Simons Fellowship in Neuroscience to RSW.

## Author Contributions

TSO and RSW conceptualized all experiments. TSO collected and analyzed all data. TSO and RSW prepared the figures and wrote the manuscript.

